# Cross-Species Variability in Lobular Geometry and Cytochrome P450 Hepatic Zonation: Insights into CYP1A2, CYP2E1, CYP2D6 and CYP3A4

**DOI:** 10.1101/2023.12.28.573567

**Authors:** Mohamed Albadry, Jonas Küttner, Jan Grzegorzewski, Olaf Dirsch, Eva Kindler, Robert Klopfleisch, Vaclav Liska, Vladimira Moulisova, Sandra Nickel, Richard Palek, Jachym Rosendorf, Sylvia Saalfeld, Utz Settmacher, Hans-Michael Tautenhahn, Matthias König, Uta Dahmen

**Author notes:** Equal contribution (Matthias König, Uta Dahmen).

## Abstract

This study explores the critical interplay between lobular geometry and the zonated distribution of cytochrome P450 (CYP) enzymes across species. We present an innovative approach to assess lobular geometry and zonation patterns using whole slide imaging (WSI). This method allows a detailed, systematic comparison of lobular structures and spatial distribution of key CYP450 enzymes and glutamine synthetase in four different species (mouse, rat, pig, and human). Our results shed light on species differences in lobular geometry and enzymatic zonation, providing critical insights for drug metabolism research. Based on our approach we could determine the minimum number of lobules required for a statistically representative analysis, an important piece of information when evaluating liver biopsies and deriving information from WSI.

## Introduction

The liver is the primary metabolic organ in all mammals. Despite the high degree of similarity between the commonly used experimental species and humans, there are still significant anatomical and functional differences that may interfere with the translation of results. With respect to liver anatomy and physiology, the following differences are evident. Humans, pigs, and mice have gallbladders, but rats lack this organ. Pigs have interlobular septa between the liver lobes, but humans, rats, and mice lack this structure. The livers of rats, mice and pigs are lobulated, whereas the human liver has no distinct lobes (Kruepunga et al., 2019).

Much less is known about functional similarity at the next smaller spatial scale. Although the microarchitecture of the liver lobule is very similar in different mammalian species, the degree of similarity in terms of lobular geometry and zonal protein expression such as cytochrome P450 (CYP) is poorly described. However, this knowledge is of great importance for the extrapolation of results from animal drug metabolism studies to the clinical situation. For example, observations made in rat hepatotoxicity studies cannot be reliably extrapolated to humans (Jaeschke et al., 2014).

Hepatic zonation refers to the spatial arrangement of metabolic processes and functions within the hepatic lobule. In the context of liver physiology, zonation describes how different zones of the liver lobule specialize in specific functions to ensure that the liver as a whole can effectively perform its diverse roles in metabolism, detoxification, bile production, and other functions. Depending on their location within the lobule, hepatocytes have different metabolic tasks. The lobule is traditionally divided into three zones: Zone 1 (periportal) - high oxygen and nutrient content, major site of oxidative processes such as gluconeogenesis and fatty acid oxidation; Zone 2 (midzonal) - mix of functions; Zone 3 (pericentral) - lower oxygen content, primary site for glycolysis, lipogenesis, and CYP-mediated drug detoxification detoxification (Ben- Moshe et al., 2019; Kietzmann, 2017; Manco & Itzkovitz, 2021). CYP isoforms involved in drug activation and steroid metabolism show a particularly pronounced zonation pattern, with high expression and selective induction in the pericentral area (Lindros, 1997).

Cytochrome P450 are key drug-metabolizing enzymes involved in the first phase of drug metabolism. They convert drugs into water-soluble products for easy elimination (Almazroo et al., 2017; Garza et al., 2022). CYPs include many isozymes present in the smooth endoplasmic reticulum and mitochondria of hepatocytes, small intestinal epithelium and proximal renal tubules (Almazroo et al., 2017). Based on the relative distribution of liver CYPs, CYP1A2, 2C8, 2C9, 2E1, 3A4 are the most abundant, whereas 2A6, 2B6, 2C19, 2D6, and 3A5 are less expressed (Paine et al., 2006; Shimada et al., 1994; Zanger & Schwab, 2013).

Few publications have addressed cross-species comparisons of CYP expression and distribution. Previous studies, such as that of Hammer and coworkers, have used targeted proteomics to obtain cross-species data on CYP protein abundance. These studies have mainly compared different in vivo and in vitro experimental systems across mice, rats and humans (Hammer et al., 2021). Hrycay reviewed the comparison and abundance of expression, function, and regulation of mouse CYP enzymes with human CYP enzymes (Hrycay & Bandiera, 2009). However, to the best of our knowledge, there has not been a comprehensive collection or comparison of histological imaging data for CYP enzymes, analyzing lobular geometry and zonal distribution pattern across different species. Such information holds significant importance in understanding the potential variations in drug metabolism and toxicity across various species, thereby facilitating the development of more precise animal models for preclinical drug evaluation.

Several methods have been explored to elucidate liver lobular geometries and their zonation patterns. For instance Voronoi theory was used to analyze liver lobular geometry in both porcine and human normal livers to provide insight into the organization of classical liver lobules as revealed by glutamine synthase (GS) staining (Lau et al., 2021). Another study applied zonal image analysis via Voronoi tessellation to discern the distribution of hypoxia markers in the hepatic lobule of steatotic livers in mice after manual annotation of central veins based on morphology on H&E and IHC for GS and liver capsule (Peleman et al., 2023). Notably, their zoned quantification of immunohistochemistry was also extended to human liver samples. Recent work introduced the tissue positioning system, a deep learning-based method that is tailored to automatically analyze zonation within the liver lobule lobule (Wang et al., 2023). All these approaches relied on manual annotation of the images, and were limited to a single or two species.

Understanding hepatic lobular geometry across species remains elusive, with little existing literature shedding light on these differences. Similarly, the zonation patterns of CYP enzymes across species are not well defined, leaving the extent of their similarities or differences largely unknown. A notable absence in the field is the lack of a mathematical framework capable of discerning geometries and zonation patterns from whole slide images, which hinders accurate and standardized analysis. Another key unresolved issue in the field is the number of lobules required for in-depth histologic analysis with direct clinical implications. Furthermore, the field of intrahepatic interlobular variability in zonal expression of CYP enzymes is underexplored despite its relevance. This is particularly important in human studies, where liver tissue is often obtained from minimally invasive needle biopsies. Given the limited nature of such samples, it’s imperative to determine the minimum number of lobules from a given species that can adequately represent the zonated expression of a given CYP enzyme.

In this study, we compared lobular geometry and CYP expression in four different species. We introduce a novel workflow to analyze lobular geometry and zonation patterns in whole slide images (WSI). Using this approach, we systematically compared the lobular geometry and zonation patterns of key CYP450 enzymes across species.

## Results

Lobular geometry, spatial distribution and zonation of CYP enzyme expression are three important features when assessing drug metabolism. We used HE-stained samples for assessing lobular geometry in four species (mouse, rat, pig, and human). Then, we visualized five enzymes (GS, CYP1A2, 2D6, 2E1, and CYP3A4) in the four species. In this study we focussed on a systematic comparison of these features in entire liver lobules following two different approaches.

We first analyzed our histological data using the classical approach consisting of a qualitative description of lobular geometry and the staining pattern followed by the statistical comparison of the extent of the surface covered by the target protein. However, this approach is not ideal to describe differences in the zonation pattern.

In a second step, we developed a novel approach focussing on the determination and comparison of the lobular geometry in all four species, followed by zonated quantification and by the calculation of the minimal number of lobules needed for the given targeted protein.

### Determination of species-specific differences in lobular geometry and spatial expression of CYP-enzymes using the classical descriptive approach

The lobular geometry was a rather robust and stable feature. In contrast, zonation patterns differed between the target proteins within a given species and within the same protein between the different species, as shown in Figure 1.

**Figure 1.**
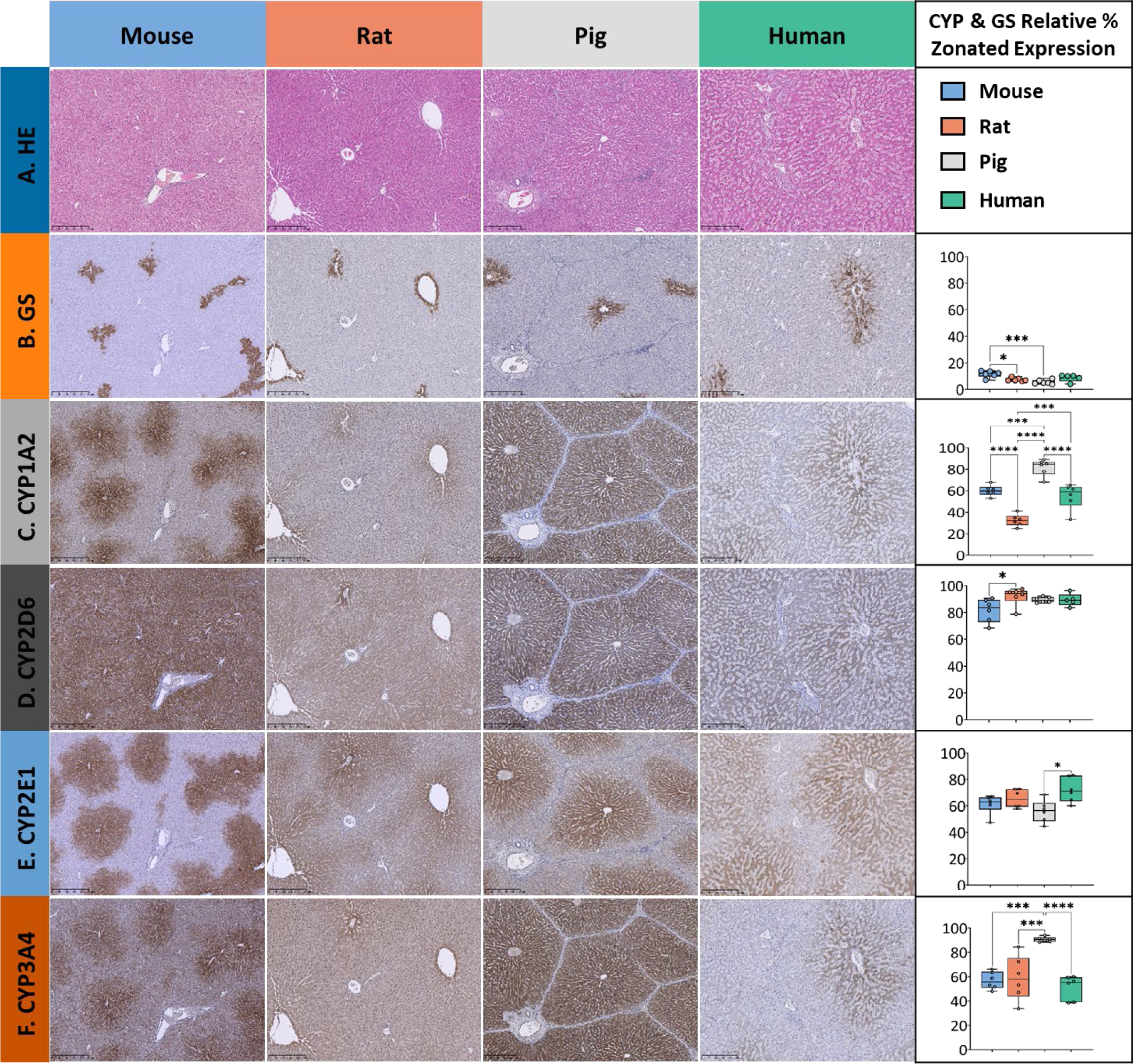
Overview of HE and extent of zonal expression as indicated by the relative surface covered by GS and CYP in liver tissue in mice, rats, pigs and humans. The different stainings are depicted in rows, with columns 1-4 corresponding to the different species, and column 5 presenting the results of the statistical analysis. (A): HE staining of normal liver tissue was used to depict lobular morphology in all species. Lobular structure appears similar except for the pig liver, where the lobules are separated by interlobular collagenous septae. (B) Glutamine synthetase (GS) staining is used to identify pericentral hepatocytes surrounding the central vein and to distinguish periportal from pericentral zones. Zonated expression is relatively similar across species, except mouse and pig liver tissue, which show a significantly different distribution. (C) CYP1A2 staining shows a similar pericentral spatial distribution and zonated expression in mice and humans, while demonstrating a significantly different zonated expression pattern in rats (pericentral) and pigs (panlobular). (D) CYP2D6 exhibits almost panlobular and similar distribution across four species. (E) CYP2E1 is observed in the pericentral region in all four species, with no substantial difference. (F) CYP3A4 shows almost identical pericentral to midzonal expression pattern in mice and humans, but panlobular in both rats and pigs. Scale bars 250 µm. Colors represent different species, with blue for mice, orange for rats, gray for pigs and green for humans. Significance levels according to descriptive one way ANOVA: * Significance level < 0.05, ** Significance level < 0.01, *** Significance level < 0.001, **** Significance level < 0.0001 (One way-ANOVA).

In terms of lobular geometry, the most striking difference between species is the presence of interlobular septa in pigs. Porcine liver lobules have septa that encircle the hepatic lobule, allowing for easy identification of the lobule. Portal fields are connected by the septa, resulting in an appearance similar to bridging fibrosis in all other species. Portal fields, even of terminal vessels, contain relatively large amounts of connective tissue.

Lobular geometry is more difficult to assess in the other three species because of the lack of clear boundaries. Lobular boundaries can be identified by the radial structure of the sinusoids. However, quantitative morphometric analysis of lobular architecture is challenging. Mice and rats are quite similar, making it difficult, even for experienced pathologists, to differentiate between the livers of the two species by morphological criteria. In contrast to pigs, the portal area of the terminal vessels lacks extracellular connective tissue and contains few histiocytes. Arteries in rodent livers are poorly visualized in contrast to those in human and porcine livers. Human lobular geometry is closer to that of rodents than to that of pigs. However, hepatic arteries are clearly visible due to the distinct appearance of the muscular vessel wall.

In terms of qualitative and quantitative assessment of CYP expression, we confirmed that all CYP enzymes are expressed in the pericentral zone of the hepatic lobules in all four species, but extend to the periportal zone to varying degrees.

GS zonated expression (Figure 1, row B) follows a similar pattern in humans, pigs, rats and mice with pericentral expression in zone 3 in the surrounding 2-3 lines of pericentral hepatocytes. Quantitative analysis of the relative area revealed a relative lobular coverage ranging from 5 to 13%. The maximum difference between species was approximately 2-fold, mouse versus pig. None of the species showed a significantly different relative coverage of GS-stained hepatocytes compared to human tissue.

CYP1A2 zonated expression (Figure 1, row C) is also most similar in human and mouse, with a strong pericentral signal extending from zone 3 into zone 2. In the rat, the signal was mostly confined to zone 3 and was of lower intensity. In pigs, the enzyme was expressed throughout the lobule and was not restricted to a specific zone. Quantitative analysis of the relative area covered by CYP1A2 showed a similar zonated expression pattern in humans and mice, with approximately 55.1 ± 11.9% and 60 ± 5.1% coverage of the hepatic lobule, respectively. In contrast, rats showed a significantly lower level of expression covering only 32.5 ± 5.5% of the lobular surface (p < 0.001), whereas pigs showed a significantly higher level of expression covering approximately 81.76 ± 7.7% of the lobular surface.

CYP2D6 zonated expression (Figure 1, row D) shows a panlobular pattern in all four species, but with homogeneous expression in humans and pigs. In rats, the expression was rather inhomogeneous, while in mice the signal was strong in the first pericentral line of hepatocytes and rather weak in the remaining zones. Quantitative analysis of the relative lobular area covered by CYP2D6 showed a similar extent of coverage, approximately 80%.

CYP2E1 zonated expression (see Figure 1, row E) follows a similar pattern and spatial distribution across species, with a strong pericentral signal extending from zone 3 into zone 2 and a weak periportal signal. Quantitative analysis of the relative area covered by CYP2E1 revealed a coverage of approximately 61.3 ± 7.2%, 65.4 ± 7%, 56 ± 8.4% of the liver in mice, rats and pigs, respectively, and approximately 72.2 ± 9.4% of the liver in humans.

CYP3A4 zonated expression (Figure 1, row F) follows a similar pattern and signal intensity in humans and mice, with a moderate to strong pericentral signal confined to zone 3 and extending into zone 2. In contrast, rat liver showed a strong signal restricted to the first line of pericentral hepatocytes, with a mild to moderate signal extending into hepatic zone 2. In pig, CYP3A4 extended throughout the lobule, covering all 3 hepatic zones. Quantitative analysis of CYP3A4 showed a consistent pattern in humans, rats and mice covering approximately 50% of the lobular surface. In contrast, porcine liver showed a significantly higher degree of zonal expression, covering approximately 90.6 ± 2% of the lobular surface with CYP3A4-positive hepatocytes.

### Novel approach: Pipeline for determination of lobular geometry, quantification and analysis of zonation patterns

We established an analysis pipeline based on whole slide images (WSI) of stained liver sections to quantify lobular geometry and zonation patterns of proteins (Figure 2, details in Method section). The pipeline consists of the following steps: (A) Registration of adjacent WSIs using Valis (Gatenbee et al., 2023), which allows the generation of multiplexed protein WSIs for the subsequent analysis. (B) Color channel separation in the immunostained WSI, providing access to the respective protein data based on color deconvolution. (C) Lobular segmentation of WSIs resulting in lobule boundaries and positional information (portality). Importantly, the presented approach does not require manual annotation of central veins and could be successfully applied to the analysis of lobular geometries and zonated quantification of proteins in mouse, rat, pig and human as well as different lobes in mouse.

**Figure 2.**
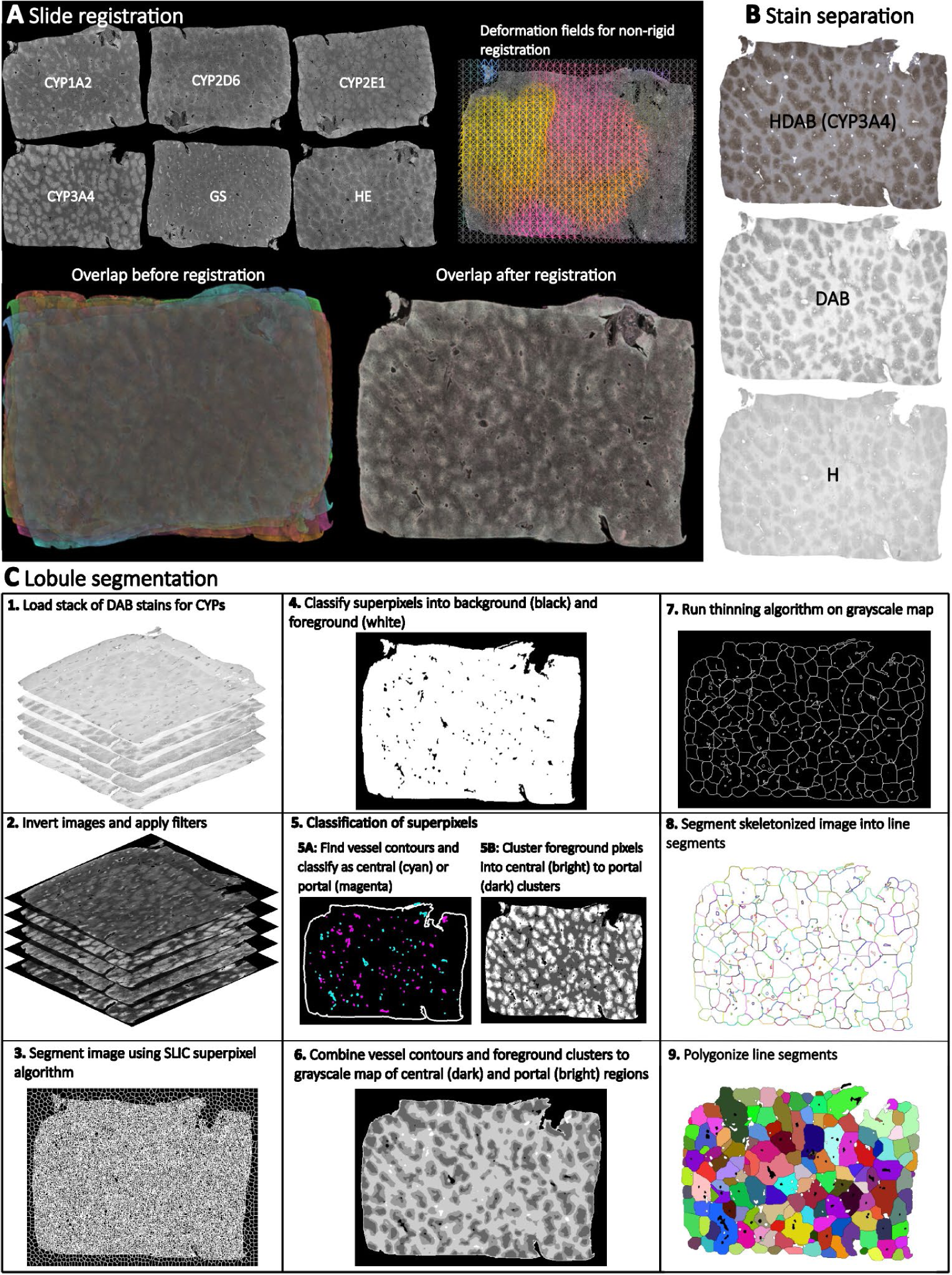
Overview of the analysis pipeline for quantifying lobular geometry and zonation patterns. The pipeline consists of the following steps: (A) Registration of HE, GS, and CYP whole slide images (WSI) using Valis, which allows the generation of multiplexed WSIs. (B) Color channel separation in the WSI. HE WSIs are separated into blue (hematoxylin) and pink (eosin), IHC WSIs are separated into blue (hematoxylin) and brown (DAB) using color deconvolution. (C) Lobular segmentation of WSIs consisting of several steps: 1. Loading a stack of DAB stains for CYPs and GS. 2. Black and white image inversion and filter application. 3. Image segmentation using SLIC (Simple Linear Iterative Clustering) superpixel algorithm to generate uniform size and regular contour superpixels. 4. Classify superpixels into background (black color) and foreground (white color). 5. Classify the superpixels: 5.A. Find vessel contours and classify vessels as central (cyan) and portal (magenta), 5.B. Cluster foreground pixels, intro central (bright) to portal (dark) clusters. 6. Combine vessel contours and foreground clusters to create a grayscale map of central (dark) and portal (light) regions. 7. Apply a thinning algorithm to the grayscale map to create a skeleton. 8. Segment the skeletonized image into line segments. 9. Polygonize the line segments to create closed polygons.

Using this approach, we systematically compared the lobular geometry and zonation patterns of key CYP450 enzymes across species (Figure 3). Lobular geometry was quantified in terms of lobular perimeter, area, compactness and minimum bounding radius followed by a statistical analysis of the differences between the species. Differences in zonation patterns in mice, rats, pigs, and humans were determined by the analysis of the zonated expression of GS, and the four CYPs (CYP1A2, 2D6, 2E1, and CYP3A4).

**Figure 3.**
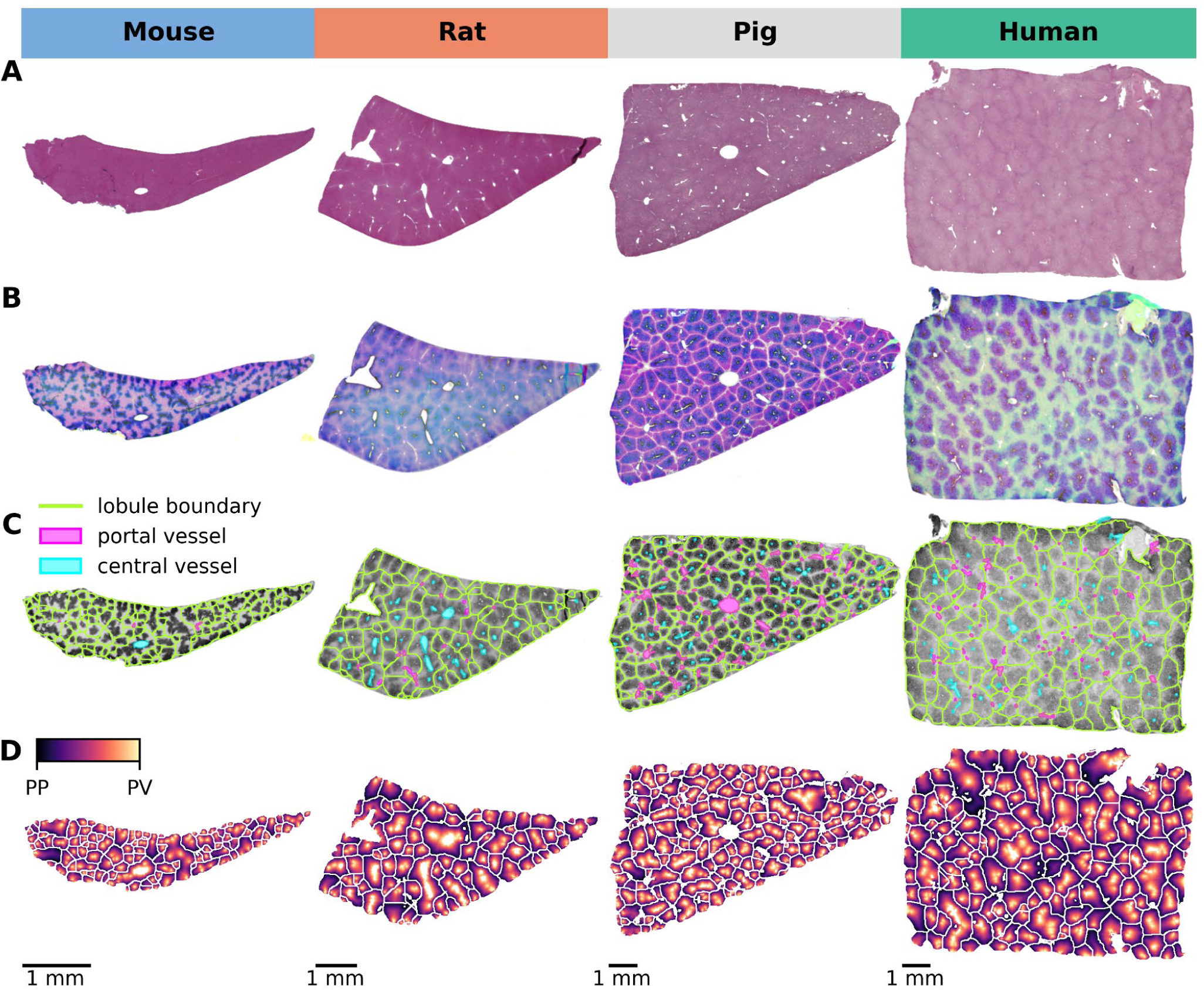
Lobular detection and position calculation. (A) HE staining of normal liver tissue from all four species showing lobular architecture. (B) Image normalization and stain separation to ensure optimal comparisons across species. (C) Detection of lobular regions providing a clear visual representation of lobular boundaries, distribution of lobules, central vessel and portal vessels on CYP2E1 staining. (D) Mapping of the central-portal distance on each lobule, allowing a quantitative analysis of the spatial arrangement of the lobules.

### Species-specific lobular geometry

Based on the segmented lobule, species-specific lobular geometry was determined using the following geometric parameters: perimeter (the measurement of the circumferential length of the outer edge of the lobule), area (surface area of the lobule), compactness (the ratio of the area of the lobule polygon to the area of a circle having the same perimeter), and minimum bounding radius (radius of the minimum bounding circle that encloses the lobule). The resulting geometric parameters for the different species and the correlation between these parameters are shown in Figure 4, with numerical values given in Supplementary Table 1-3.

**Figure 4.**
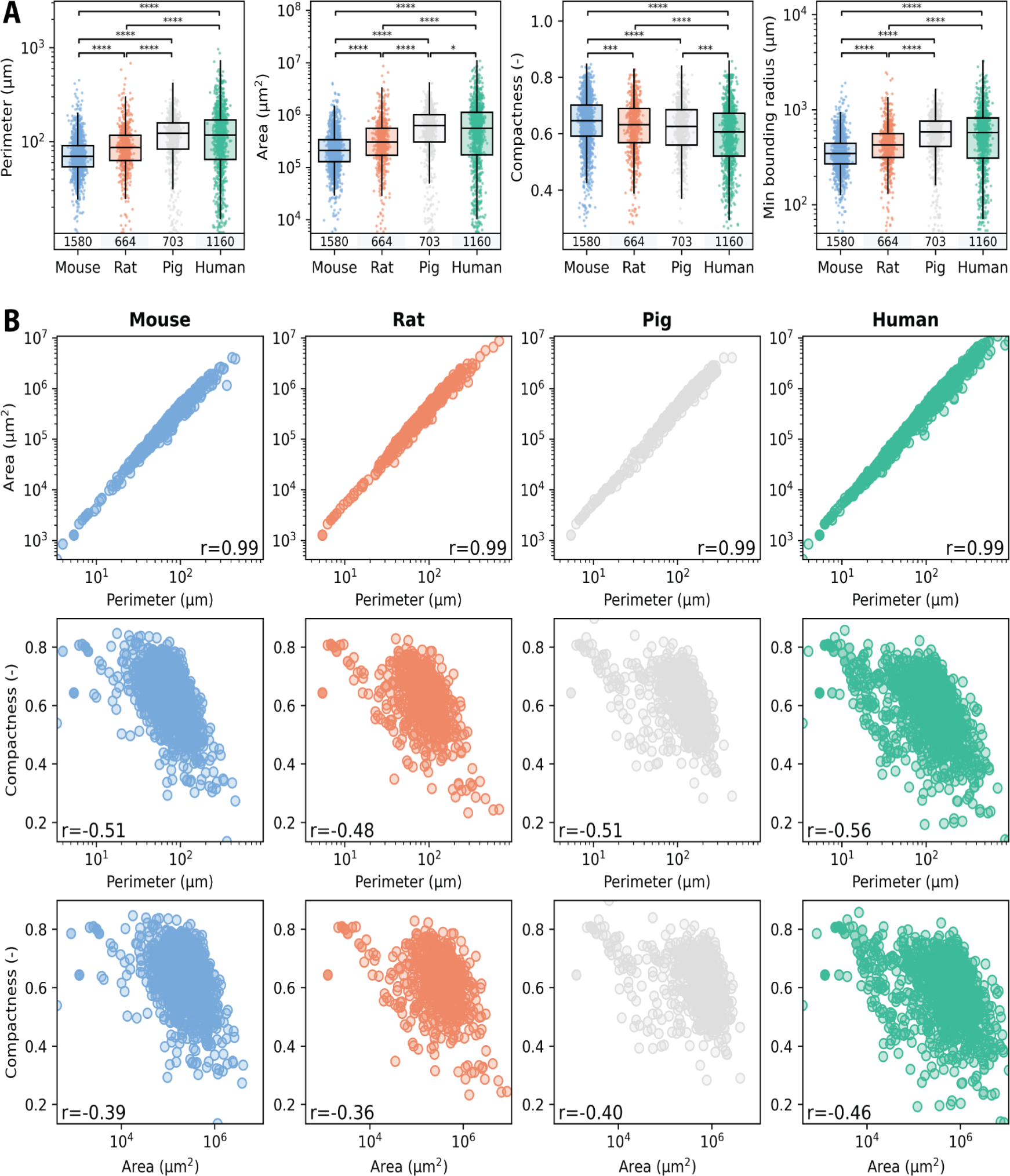
Comparison of lobular geometry in the four different species. (A) Quantification of species-specific lobular geometry. For all species the lobular perimeter, area, compactness, and the minimum bounding radius was calculated for all lobuli per section. Box blots and point clouds of these parameters are depicted. Boxes represent quartiles Q1 and Q3. The upper whisker and lower whisker extend to last datum less than Q3 + 1.5 * IQR, and the first datum greater than Q1 - 1.5 * IQR, respectively. IQR denotes inter quartile range (Q3-Q1). Significance levels: * p<0.05, ** p<0.01, *** p<0.001, **** p<0.0001. (B) Correlation between the geometric parameterswas assessed using the Spearman rank correlation coefficient. Colors represent different species, with blue for mice, orange for rats, gray for pigs and green for humans.

Geometric parameters were calculated for 1580 segmented lobules in mice, 664 in rats, 703 in pigs and 1160 in humans. Lobule size increases slightly from mouse, rat, pig to human, i.e., the larger the species the larger the lobule. The mean ± SD perimeter of a liver lobule increased from mouse 76.1 ± 39.6 µm, rat 96.1 ± 62.4 µm, pig 121.1 ± 60.6 µm to human 130.9 ± 98.9 µm with an area of mouse 282 730 ± 292549 µm^2^, rat 470198 ± 646351 µm^2^, pig 714159 ± 565512 µm^2^ to human 879809 ± 1159576 µm^2^ and a minimum bounding radius of 372.6 ± 178.6 µm, rat 463.8 ± 270.2 µm, pig 580.4 ± 276.0 µm to human 615.1 ± 419.9 µm. In contrast, the compactness, which is a parameter of the roundness of the given lobule, decreased with increasing species size and was highest in mice 0.64 ± 0.09, followed by rats 0.62 ± 0.10, pigs 0.62 ± 0.09, and humans 0.59 ± 0.12. Statistical comparison of the mediansrevealed significant but rather small differences of almost all parameters between the species. A similar large individual variability was observed in all species in the geometric parameters, with slightly larger variability in humans compared to the other species. Results of the geometric parameters from different subjects were highly similar (Supplementary Figure 1), i.e., no intra-individual variability in the geometric parameters could be observed. In addition, for mice, sections for the different liver lobes were available and could be compared (Supplementary Figure 2). Determination of the geometric parameters for the different lobes resulted in comparable geometric parameters, i.e., no differences could be observed between left lateral lobe (LLL), median lobe (ML), right lobe (RL), and caudate lobe (CL). I.e., neither intra-lobe variability in mice nor intra-subject variability in mice, rats, pigs and humans was observed in the geometric parameters, but significant differences between species.

A high variability in geometric parameters between different lobules was observed, as evident by the large SDs and interquartile ranges in all four species. One explanation could be the varying position, size and the 3-D shape of the lobule in respect to the 2-D sectioning plane.

The correlation structure for the different geometric parameters was very similar between the different species (Figure 4B). Area and perimeter showed very strong positive correlation (r=0.99) in all species. As expected, hepatic lobules with a larger area also have a larger perimeter. In contrast, compactness showed a weak to moderate negative correlation to perimeter (r in [-0.48, -0.56]) and area (r in [-0.36, -0.46]), i.e. the larger the lobule, the less compact they are possibly due to variable sectioning angles leading to a rather ovaloid appearance of the large lobule.

The similarities in the median and range of the geometric parameters across the species (Figure 4A) and the very similar correlation structure between species in the geometric parameters (Figure 4B) implies that the underlying complex 3- dimensional structure of the hepatic lobule must be rather similar. In conclusion, lobular geometry seems to be a robust feature, with little interindividual and even little interspecies variability, but large variability between different lobules.

### Quantification of zonated expression from whole slide images: Gradients & zonation patterns

The subsequent analysis quantified the zonated expression of CYP enzymes along the portal-venous axis in the entire liver lobules of the four species (Figure 5). Based on the calculated positions within the segmented lobules, we determined the position-dependent protein expression. Specifically, we assigned positions to all pixels within a lobule, ranging from periportal (0) to perivenous (1), based on their proximity to the nearest periportal or perivenous region. Using these positions, we determined the zonation patterns of GS and CYP proteins across lobules and species. Analysis of the combined zonated expression of all markers revealed distinct patterns for the proteins and species.

**Figure 5.**
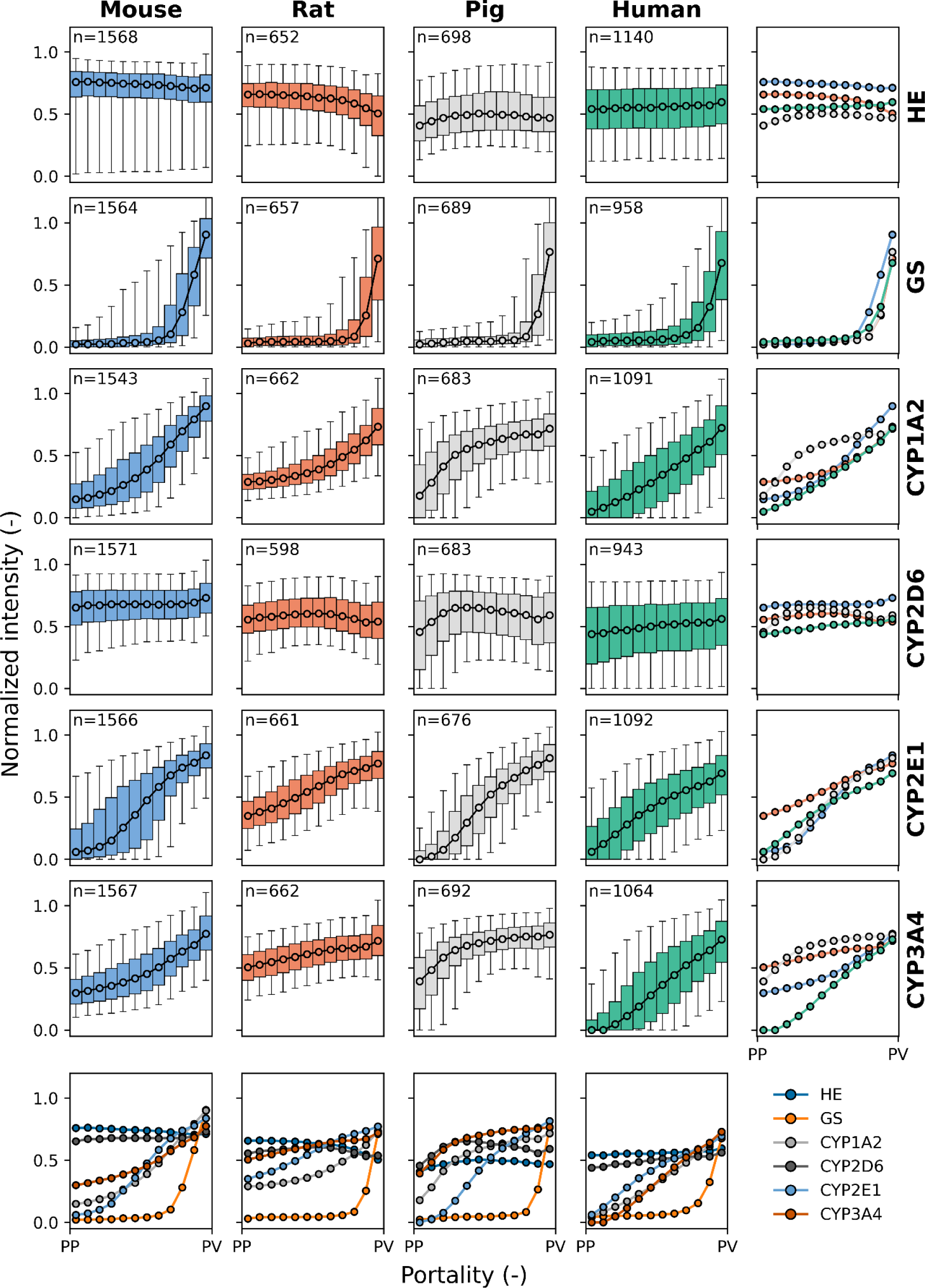
Species comparison of protein zonation. Zonation patterns of HE, GS, CYP1A2, CYP2D6, CYP2E1, CYP3A4 in mouse (blue), rat (orange), pig (gray) and human (green). Normalized protein intensity (per slide) is plotted against portality (relative position between periportal (PP) and perivenous (PV) zones in each lobule). Data were binned in 12 bins from PP to PV. Median values are shown for all lobuli of all individuals. Box plots correspond to median, interquartile range with whiskers at 5%, 95% percentile. Colors represent different species: blue for mice, orange for rats, gray for pigs, and green for humans. n: number of lobuli for the respective analysis.

HE-staining appeared as a consistent flat line across all species. This result was expected as HE-staining does not indicate protein expression along the sinusoid but rather delineates the morphological structure of the hepatic lobule and no zonation differences are expected.

GS showed a similar gradient and zonation pattern along the entire portal-venous axis in the four species, as shown by the superimposed plots of normalized staining intensity. GS was predominantly localized in zone 3, encompassing the 2- 3 lines of pericentral hepatocytes, with no periportal distribution pattern in the four species.

CYP1A2 showed similar gradient and zonation patterns in mice, rats and humans, predominantly located in zone 3 and extending into zone 2 within the adjacent 5-6 rows of pericentral hepatocytes. In pigs, however, the gradient distribution of normalized intensity was mainly seen in zones 3 and 2 and extended into zone 1 of the periportal hepatocytes.

CYP2D6 showed a constant flat zonation pattern similar to HE in all four species, with a more panlobular distribution within the lobule along the portal-venous axis. CYP2D6 was the only CYP analyzed that did not show a clear zonation pattern with higher protein content in the perivenous region compared to the periportal region.

CYP2E1 showed a linear gradient distribution of normalized intensity throughout the liver lobules in different species, predominantly in zones 3 and 2. In rats, there was a higher intensity in zone 1, with a flatter gradient than in the other species.

The intensity of the CYP3A4 gradient was similar in mice and humans, mainly in zone 3 and extending into zone 2. Conversely, rats and pigs exhibited a similar but distinct gradient distribution to mice and humans across the liver lobules, with CYP3A4 normalized intensity higher in zones 3 and 2 and extending to periportal hepatocytes in zone 1. The strongest periportal to perivenous gradient was observed in humans.

This innovative approach to zonation assessment does allow comparative analysis of the zonation patterns of multiple target proteins, such as the four different CYP proteins, in a single species as well as the comparison of single proteins between species. The superimposed curves clearly visualize that the zonated expression patterns in mice and humans are more similar, while those in rats and pigs are more different.

### Number of lobules required to determine parameters

Next, we determined the number of lobules required for a representative analysis of geometric parameters, zonation patterns, and relative expression (Figure 6).

**Figure 6.**
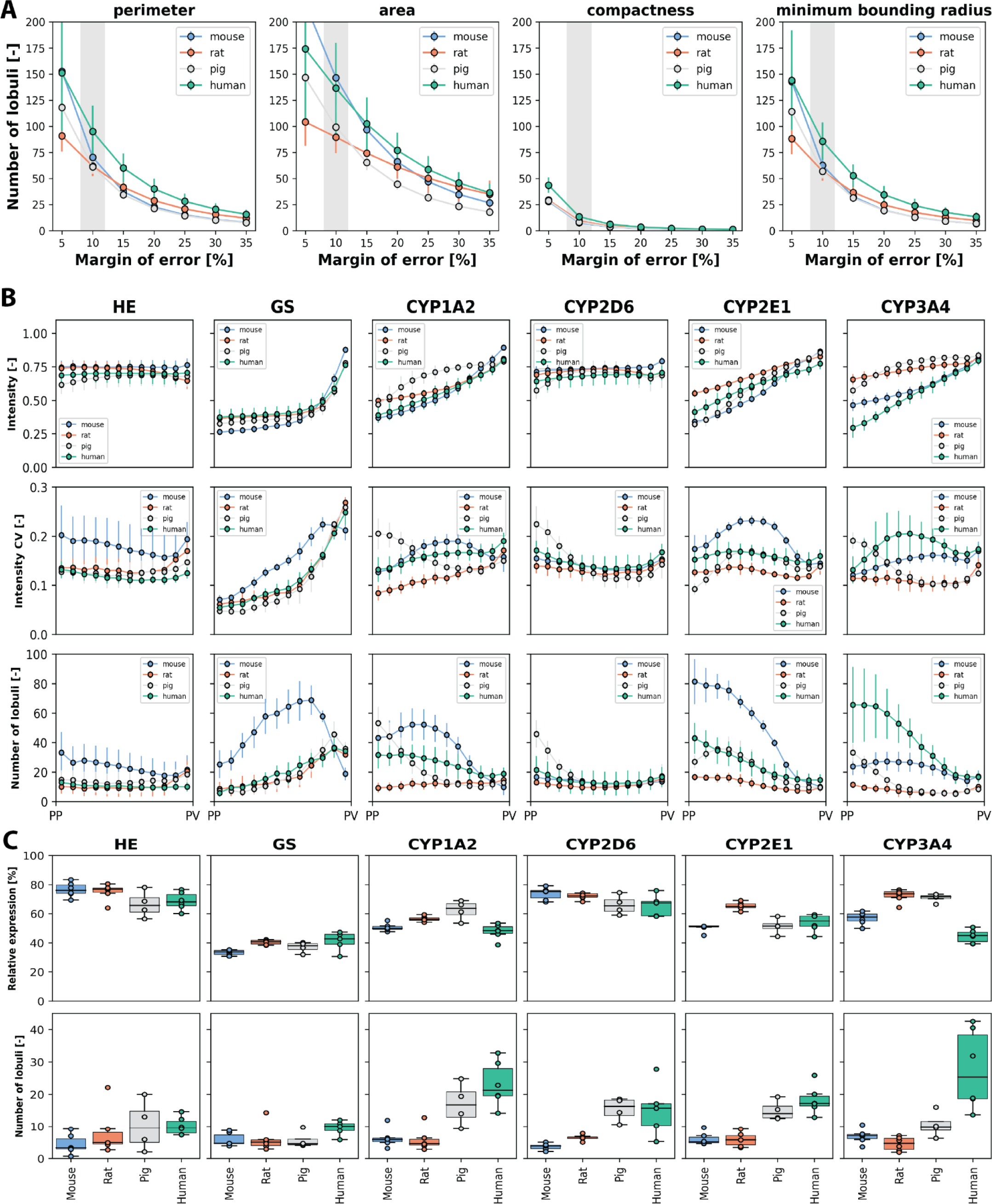
Number of lobules required to determine geometric parameters, zonation patterns, and relative expression. Species are color coded as mouse (blue), rat (orange), pig (gray) and human (green). (A) Number of required lobules to determine the geometric parameters area, perimeter, compactness, and minimum boundary radius for a given margin of error (5% to 35%) and a confidence level of 95%. Data expressed as mean ± SD for the subjects. (B) Number of required lobules to determine the protein zonation pattern, i.e., the normalized intensity for the 12 bins with a confidence level of 95% and a margin of error of 10%. Depicted are normalized intensity mean ± SD for the subjects (top), the coefficient of variation of the normalized intensity as mean ± SD for the subjects (middle) and the required number of lobules for estimation of the zonation pattern as mean ± SD for the subjects (bottom). (C) Number of required lobules to determine the relative protein expression per lobulus with a confidence level of 95% and a margin of error of 10%. Relative expression as mean values for individual subjects (averaged over all lobuli per subject) and the corresponding boxplot of the means (top). Number of lobules required for individual subjects and the corresponding boxplot (bottom).

First, we determined the number of lobules required to calculate mean geometric parameters with 95% confidence at a given margin of error (Figure 6A, data in Supplementary Table 2 and 3). For example, To calculate compactness with 95% confidence and a 10% margin of error, n = 7.7 ± 1.0 lobules are required in mice, 9.3 ± 1.1 in rats, 8.2 ± 1.0 in pigs, and 13.4 ± 1.9 in humans.

Key findings included: (i) A larger margin of error reduces the number of lobules required for analysis. The dependence of the number of lobules required on the margin of error is highly non-linear, i.e. a small relaxation of the possible margin of error requires much fewer lobules. (ii) Human geometric parameters require a larger number of lobules for accurate determination compared to mice. The number of lobules required for rats and pigs is between those required for humans and mice. (iii) Different geometric parameters require different numbers of lobules for accurate evaluation. For example, area requires more lobules than perimeter and minimum boundary radius, while compactness requires the least number of lobules for accurate quantification. (iv) There is significant variability between biopsies of the same species. This variability suggests that there is also some intrahepatic variability in terms of geometric parameters, resulting in slightly different numbers of lobules required for a given level of targeting accuracy in a given biopsy.

We then determined the number of lobules required to calculate zonation patterns with 95% confidence and a 10% margin of error (Figure 6B). The number of lobules required for the zonation pattern varied with the protein of interest as well as the species. For many positions and proteins, < 20 lobules were sufficient to determine the mean expression for a given zonal position. Determination of zonated protein expression of GS and CYP2E1 in mice required much more lobules than in the other species with up to 60 lobules for GS and 80 for CYP2E1. CYP3A4 in humans required up to 60 lobules for periportal protein levels. An important factor for the required number of lobules was the coefficient of variance of the intensity at the given position. A large variability in protein expression at a given position resulted in a much higher number of lobules for a reliable determination of the mean intensity. The protein pattern in a liver lobule is very similar for most proteins, but the variability can be very different between species.

Finally, we determined the number of lobules required to determine the relative expression of proteins within a lobule with 95% confidence and 10% error (Figure 6C, data in Supplementary Table 4). The number of lobules required for HE, and GS staining was almost similar across different species with up to 10 lobules in humans. Determination of the relative expression of different CYP proteins in mice and rats was relatively similar and required less lobules than in other species with up to 23 lobules for CYP1A2, 15 for CYP2D6, 18 for CYP2E1, and 28 for CYP3A4 in humans.

An important finding is that significantly fewer lobules are required to determine the relative expression of a protein in a lobule than to determine the lobular pattern. To determine relative protein expression, the highest number of lobules was required for human liver tissue, followed by pig, rat and mouse liver tissue.

## Discussion

Within this work we present a novel approach to assess lobular geometry and zonation patterns using whole slide imaging (WSI) without the need for manual image annotation. This method allowed a detailed, systematic comparison of lobular structures and the spatial distribution of key CYP450 enzymes in four different species (mice, rats, pigs and humans). Importantly, our approach does not require time-consuming manual annotation of the WSI by experts in contrast to other published methods segmenting the hepatic lobule (Lau et al., 2021; Peleman et al., 2023; Schwen et al., 2016; Wang et al., 2023).

### Lobular geometry

We showed here for the first time that lobular geometry is a rather robust and stable parameter with little differences within the liver, i.e. between liver lobes, between subjects and also between species. The area and perimeter of the lobule increases slightly with increasing species size from mouse, to rat, to pig, to human. The main geometrical difference between the livers of the different species is within the lobar structure with rodent and pig liver being lobulated whereas the human liver is rather compact (Kruepunga et al., 2019).

Furthermore, the absolute size/volume of the liver varies, with adult human livers weighing approximately 1500 g (constituting 2 to 3% of total body weight). Rodent liver weight, on the other hand, ranges from 2 to 3 g, equivalent to 3 to 5% of body weight in mice and 4 to 5 g, equivalent to 2 to 3% of body weight in rats (Rogers & Dintzis, 2018). Pigs’ liver weight ranges from 1000 to 2000 g, equivalent to 1.4 to 2% of body weight (Elefson et al., 2021). Since the volume is different, the absolute number of hepatic lobules is highly different. However, the hepatic microarchitecture is surprisingly robust, with significant but rather small differences in the related 2D geometrical parameters. This finding has important implications for computational modeling of the liver, which allows reusing lobular mathematical models of the liver across different species for simulation purposes with minor modifications (Lambers et al., 2023; Ricken et al., 2015).

Lobule area and perimeter were highly correlated (correlation coefficient r = 0.99), whereas compactness and area respectively compactness and perimeter were weakly negatively correlated (r < 0.5). This finding suggests that the 3- dimensional structure of a hepatic lobule resembles possibly a more or less asymmetric ovaloid sphere. Here more technical work is required to not only register a stack of stained sections to extract the 2D lobular shape, but to go for a 3D reconstruction of entire hepatic lobules to determine the 3D structure and packaging of the liver lobules in the context of the vascular tree. Reconstruction of the entire hepatic lobule and the vascular tree and sinusoidal network pose major technical challenges beyond our approach.

As part of the work we developed a robust method allowing non-rigid registration of a small stack of WSI sections resulting in multiplex protein images. While some spatial resolution is lost in this approach due to the registration, the results are more then sufficient to determine lobular geometries and zonation patterns of proteins, but not sufficient to resolve subcellular information. Importantly, the presented workflow is straight-forward applicable to immunofluorescence WSI images with multiple proteins which would enable it to determine zonation patterns and lobular geometries with high-spatial resolution.

The developed image processing workflow is a crucial prerequisite for further related imaging and modeling studies. For instance, the established automatic registration and lobule segmentation without prior annotation is an important step to extend the analysis for larger datasets as required for 3D reconstruction of the hepatic lobule and extraction of the vascular and sinusoidal network. Both again is the absolute requirement for flow simulation using real lobular geometry which in turn is needed for further simulation of the perfusion-function relationship.

### Cyp expression

Drug metabolism studies have been performed for many years, both in humans and experimental animals involving different species. The underlying assumption is that the results obtained in animal studies are reflecting the clinical situation.

So far, not much attention was paid to spatial and especially zonal distribution of CYP expression in different species, although it might be of high relevance to explain differences in drug metabolism and toxicity between species. Within this work we performed a systematic analysis of the spatial distribution and zonation patterns of four CYP450 enzymes (CYP1A2, CYP2D6, CYP2E1 and CYP3A4) in four species (mice, rats, pigs and humans). Interspecies comparison of CYPs is very limited in the literature, with one study comparing the panlobular expression of CYP3A4 in livers of adult minipigs with the reported pericentral to midzonal expression observed in humans (Van Peer et al., 2014).

It is highly likely that the extent and zonated distribution of CYP-expression are affecting metabolic activity and might be decisive for the interpretation of translational studies. We observed high similarities in zonal GS expression as well as in CYP2E1 expression between species. Our observation is in line with others (Martignoni et al., 2006) reporting also a high degree of similarity in terms of catalytic activity. The authors concluded that CYP2E1 activity did not show large differences between species, and extrapolation between species appeared to hold quite well. In contrast, the species-specific isoforms of CYP1A, -2C, -2D and -3A show considerable interspecies differences in terms of expression pattern, as observed in our study, but also in catalytic activity, as reported by (Dalgaard, 2015; Martignoni et al., 2006). These differences in the extent of expression and the catalytic activity could be the reason for striking differences in terms of hepatotoxicity.

Knowing the species-specific features of drug metabolism such as differences and similarities in CYP patterns, will enable us to better predict therapeutic efficacy and toxicity according to species, as well as generating safer and more effective therapeutic plans. It may also have a substantial effect on drug testing and preclinical drug development.

### Minimal number of lobules

In clinical pathology, it is highly recommended to assess a minimum number of lobules to obtain a reliable pathological diagnosis. Misdiagnosis can happen as a result of limited sample size, particularly when the recommended minimal number of portal fields are not obtained, the disease process is localized, and the interpreter lacks experience. The optimal length for a liver biopsy is reported to be 1-4 cm and weighs 10-50 mg (Sherlock & Dooley, 2002). The majority of hepatopathologists are satisfied when given a biopsy specimen that includes at least six to eight portal triads, especially in cases of chronic liver disease where the extent of damage may vary among portal triads (Bravo et al., 2001). Agarwal (2022) reported recently that a clinical liver biopsy should contain at least 10 complete portal fields to reliably diagnose allograft rejection (Agarwal et al., 2022). This number is purely based on pathological experience, but not on detailed quantification of lobular geometry and quantification of staining patterns.

Within this work we determined the minimal number of lobules required to determine geometrical parameters such as area and perimeter, protein zonation patterns, and percentages of stained area. Importantly, the minimal number of lobules required for a reliable quantitative analysis of lobular geometry varied for the given parameter and the allowed margin of error for the estimate. The most robust parameter was the compactness requiring 6.5 - 16.2 lobules, whereas the area was less robust, requiring as many as 70 - 230 lobules for a confidence of 95% at a margin of error of 10%. The relatively large number of lobules required for a reliable estimate of the lobular geometry is a consequence of the large variability in the geometric parameters between different lobules from the same subject, whereas the means between individual or even species are rather similar.

Calculation of the minimum number of lobules needed to assess the zonated distribution showed large differences between species. The main factor contributing to a large number of required lobules was a large coefficient of variation in the protein amount for the spatial locations, i.e. the more heterogeneous the zonation patterns are between different lobules on a sample, the higher the number of lobules needed for a reliable estimate. These differences in zonation patterns between different lobules could have important implications for toxicity or spatial drug metabolism.

When performing the same calculation for CYP staining in terms of relative expression covered by a given staining, much fewer lobules were required to achieve the same level of confidence, with < 10 lobuli required for rat and mouse, 10 - 20 lobules for pig, and 10 - 30 lobules for human, depending on the protein of interest.

However, in this study, we only looked at normal livers in the four species. It is highly likely that the minimal number of lobules will increase in case of structural abnormalities. A general pattern observed was that the higher the variability in the readout of interest between different lobules, the more lobules are required for a reliable estimate of the parameter.

Even for simple parameters such as area, a large number of lobules are required which can be obtained from WSI of a large liver sample, but not from a single liver core biopsy. Many parameters used in histopathological scoring systems show a large spatial variability representing a challenge for reliable assessment.

Taken together, the minimal number of lobules depends substantially on the species, the parameter and probably also the morphology of the liver. For many parameters, this number is much higher compared to the pathological standards for clinical diagnosis. Our results suggest that a quantitative analysis of clinical biopsies should be interpreted with caution. This may have implications for clinical pathology, where the number of lobules determined here can never be reached with a single biopsy.

## Conclusion

In conclusion, we presented a novel approach to assess lobular geometry and zonation patterns using whole slide imaging (WSI) without the need for manual image annotation. This method allowed a detailed, systematic comparison of lobular structures and the spatial distribution of key CYP enzymes (CYP1A2, CYP2D6, CYP2E1, and CYP3A4) and GS in different species (mice, rats, pigs, and humans). The approach allowed to determine the minimum number of lobules required for statistically representative analysis, an important piece of information when evaluating liver biopsies.

## Material and methods

### Samples

Archived formalin-fixed and paraffin-embedded liver samples from normal rats, mice, pigs and humans obtained from other studies were used. All experiments and housing of the animals were performed according to current German regulations and guidelines for animal welfare and the ARRIVE Guidelines for Reporting Animal Research. Approval for the mouse study was given by the Thüringer Landesamt für Verbraucherschutz, Thuringia (Approval-Number: UKJ-19-020, see also (Albadry et al., 2022), for the rat study Reg.-Nr.02-043-10, Approval for the pig study was given by the Commission of Work with Experimental Animals under the Czech Republic’s Ministry of Agriculture, project ID: MSMT-15629/2020-4.

Human samples were collected during clinically indicated liver surgeries in the time period 2019 as approved by the ethical vote 2018-1246-Material. Normal samples without additional pathologies were selected by subjecting whole slides scans of the HE-stained sections to the independent assessment by four scientists (UD; MA; HT, OD) one being a board certified pathologist.

### HE-staining and IHC

Paraffin-embedded liver tissue samples were subjected to hematoxylin-eosin (HE)-staining after cutting 3µm sections to assess lobular geometry. HE-staining was initiated by deparaffinization and rehydration using descendant grades of alcohol. Sections were immersed in hematoxylin and eosin solution, followed by dehydration using ascendant grades of alcohol. Slides were mounted and digitized to a digital format using a whole slide scanner (L11600, Hamamatsu, Japan) that was fitted with the NDP.view2Plus Image viewing software (Version U12388-02) at 40 × magnification.

Immunohistochemistry was employed in order to evaluate and quantify the spatial distribution of glutamine synthetase (GS) and four different cytochrome P450 (CYP) enzymes as mentioned before (Albadry et al., 2022).

The staining procedure was conducted utilizing a series of consecutive sections, each measuring 3 µm in thickness, derived from liver tissue that had been fixed in formalin and embedded in paraffin. Five different antibodies were utilized for the visualization of GS, CYP1A2, CYP2D6, CYP2E1, and CYP3A4 see Supplementary Table 5. Following the process of deparaffinization and rehydration using descending grades of ethanol, heat-based epitope retrieval was initiated. This involved the use of Trisodium-citrate buffer with a pH of 6.1 and a steamer set at a temperature of 100°C for a duration of 30 minutes. Subsequently, the samples were allowed to cool down at room temperature for a period of 20 minutes. The activity of endogenous tissue peroxidase was inhibited by treatment with a 3% hydrogen peroxide solution. The endogenous IgG was effectively inhibited using a commercially available protein block (ab64226, Abcam, Germany). The subsequent stage involved an overnight incubation at a temperature of 4°C, utilizing the corresponding CYP antibody as indicated in Table 2. Rabbit-specific HRP/DAB IHC detection system (ab236469, Abcam) was utilized for Rabbit polyclonal primary antibodies (CYP2D6, 2E1, and 3A4) for 40 minutes at room temperature. For mouse monoclonal primary antibodies (GS, and CYP1A2), additional biotinylation of the primary antibody was performed using Dako Animal Research Kit Peroxidase (K3954, Dako, Denmark). Additionally, endogenous avidin and biotin activity was inhibited using Avidin/Biotin Blocking kit (ab64212, Abcam), followed by application of the Avidin-HRP complex. DAB-chromogen was implemented for 3–5 minutes to visualize the reaction. Counterstaining was performed using Dako hematoxylin (CS700, Dako, Denmark) for 6–8 minutes. During each staining trial, a single slide was designated for the negative reagent-control condition, wherein the primary antibody was intentionally omitted. Slides were mounted and digitized to a digital format using a whole slide scanner (L11600, Hamamatsu, Japan) that was fitted with the NDP.view2Plus Image viewing software (Version U12388-02) at 40 × magnification.

### Qualitative assessment of zonated CYP-expression

CYP-expression was described qualitatively by taking the extent and zonal distribution (periportal, midzonal, and pericentral) and the signal intensity (mild, moderate, or strong) into account.

### Image analysis based quantification of spatial distribution of CYP-expression

The quantification of zonated CYP spatial distribution was performed using Histokat, a proprietary software developed by Fraunhofer MEVIS that utilizes a machine-learning algorithm. This approach segments the entirety of the slide scan into discrete square tiles of a predetermined size. The software was trained to identify certain features or patterns by utilizing a minimum of 30 tiles per image from distinct liver lobes within a single slide. This training involved the use of various representative images from the series.

Spatial distribution of CYP-expression, expressed as relative surface covered by CYP-stained hepatocytes, was quantified using pattern recognition algorithm (generic classification 128) as mentioned before (Albadry et al., 2022).

### Statistical analysis

For analysis of CYP zonated expression across four different species (mice, rats, pigs, and humans), a descriptive ordinary one-way analysis of variance (ANOVA) was performed. The Tukey’s multiple comparisons test was conducted using GraphPad Prism version 9.3.1(471) for Windows, a software developed by GraphPad Software, San Diego, California, USA, www.graphpad.com. The data were presented as the mean value ± standard deviation. Statistical significance was determined for differences when p-values were below 0.05.

### Pipeline for quantification of lobular geometries and zonation patterns

The image analysis pipeline consisted of the following main steps: region of interest (ROI) detection, image registration, stain separation, and liver lobule segmentation. The WSI images were provided as RGB images in NDPI format. For each subject, HE and HDAB stained images were provided for the proteins GS, CYP1A2, CYP2D6, GYP2E1, and CYP3A4. All images from a single subject were from adjacent slides.

### ROI detection

First, the source image files provided in NDPI format were annotated using QPath to mark the tissue samples on the slides. Subsequently, the whole slide images were loaded on the lowest resolution level with openslide (Goode et al., 2013), converted to grayscale, and thresholded using a binary threshold to get a foreground mask marking the tissue portion of the slide. Contour detection was utilized to transform the binary mask to polygons. The polygons were checked and selected if they contained the coordinates of the tissue annotation. The bounding boxes of the selected polygons were then used to load the region of interest (ROI) of liver tissue of the source image in full resolution. The ROIs were saved as OME-TIFF. For mice subjects, 4 ROIs from different liver lobes were present on the slide. After ROI detection, the ROIs were matched based on the similarity of the tissue mask obtained by OTSU thresholding. In case of very similar shaped lobes, the matching failed and the ROIS were mapped by hand. For contour detection and thresholding, OpenCV (Bradski & Kaehler, 2008) implementations of the respective algorithms were used.

### Image registration

The set of images for each ROI was registered using VALIS, virtual alignment of pathology image series for multi-gigapixel whole slide images (Gatenbee et al., 2023), with default settings, and the registered ROIs were saved as OME-TIFF.

### Stain separation

Registration was followed by stain separation, resulting in two single-channel images of the H-stain and the DAB or E- stain, respectively. The resulting images were stored in the ZARR format [https://zarr.readthedocs.io/en/stable/] with eight pyramidal layers. For stain separation, we used a color deconvolution algorithm introduced by Macenko (Macenko et al., 2009).

### Lobule segmentation

We used a classical image analysis approach to segment the lobule boundaries. The following procedure was applied to the image set for each subject and ROI of the dataset. (1) The DAB stain for the image set was loaded for the proteins GS, CYP1A2, CYP2D6, GYP2E1, CYP3A4 at resolution level 5 (2.5x magnification). The registered protein ROIs were stacked into a 5-channel image. Protein images where the number of foreground pixels for a protein image was less than 80% of the median number of foreground pixels across all protein images were discarded. (2) Protein images were inverted so that bright regions correspond to high absorbance (high expression). Pixels where one of the channels was zero (i.e. background) were set to zero. Image filters were then applied to each channel. First, a median filter was applied, and the image was convolved to resolution level six. Adaptive histogram normalization was applied to reduce global differences in illumination and staining. Finally, after applying a median filter, the image was convolved to resolution level seven and each channel was normalized to the maximum intensity of the channel. (3) The 5-channel images were segmented using OpenCV’s superpixelization implementation, which assimilates similar pixels into a larger superpixel. (4) The superpixels were divided into foreground and background pixels. Pixels were classified as background if more than 10% of the pixels in a superpixel were zero. (5) Foreground pixels were grouped into three zones: pericentral, midzone, periportal.

Therefore, each superpixel was transformed into a 5-channel vector containing the mean intensity for the channel in the superpixel. The vectors were clustered into three clusters using the K-means algorithm implementation of scikit-learn (Pedregosa et al., 2011). The clusters were sorted by the Euclidean distance of the cluster centers. The highest distance corresponded to the pericentral zone, since expression is high for all proteins in this zone. The labels were mapped back to the superpixel representation, marking each foreground superpixel as either pericentral, periportal, or midzone.

A mask was created from the background pixels and contour detection was applied, yielding the contours of vessels and the tissue boundary. The contours were then classified into pericentral and periportal vessels. This was done by analyzing the foreground pixels in the adjacent area around the vessel. The occurrence of each label (pericentral, midzone, periportal) was counted. It was expected that the number of pericentral labels would be higher for pericentral vessels and vice versa for periportal vessels. Based on this reasoning, the count vectors for all vessels were clustered into two clusters using the K-means algorithm.

(6) A grayscale representation was created from the clustered foreground and vessel contours, with pericentral and periportal vessels colored black (zero) and white (255), respectively. The foreground zones were uniformly distributed across the grayscale spectrum in between, with dark and light corresponding to pericentral and periportal zones, respectively. (7) An OpenCV implementation of a thinning algorithm was used to skeletonize the grayscale representation. The remaining lines are located in the expected center of the periportal zones and vessels (bright zones), marking the potential boundaries. (8) To polygonize the skeletonized image, we extracted line segments that we could polygonize. A line segment consists of adjacent pixels, where each pixel has at max two neighbors in the perpendicular or diagonally connected pixels. We implemented a pixel walking algorithm that yields line segments including the connecting pixels. The algorithm extends a line segment recursively. For a given line segment, it analyzes the neighbors of the last appended pixel. If only one neighbor is found, it is appended to the segment and the process is repeated. If no neighbor is found, the segment is terminated. If multiple neighbors are found, the segment is terminated and new segments are created for each neighbor. When a segment is finished, the next initial segment is taken from the list of initial segments. The process is repeated until there are no initial segments left. (9) Finally, we used the shapely (Gillies et al., 2023) library to polygonize the set of line segments we obtained in the previous steps. All line segments that are not part of a closing circle are discarded.

### Generating the Portality Map

To analyze expression gradients, we calculated the relative position of each pixel in a lobule. The portality p was defined as

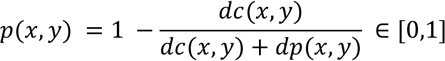

where dc and dp are the distance of a position to the nearest central and portal pixel, respectively. To do this, we used the lobule boundary polygon and the vessel polygons to create a periportal and a pericentral mask. For these masks, the distance transformation was calculated using the OpenCV implementation. This function calculates the distance to the nearest background pixel, resulting in two maps of the central and portal distance for each pixel in the lobule. For each protein, the intensity was background corrected and normalized by

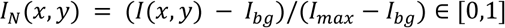

where I(x,y), Ibg, Imax denote the intensity of the pixel, the background intensity, and the maximum intensity of the slide, respectively. The maximum intensity was defined as the 99% quantile of the foreground pixels. The background intensity was estimated by the 20% percentile of the foreground pixels of the GS slide for each subject. GS expression covers only a very small area of the lobule leaving the remainder as a robust background estimate. For each pixel, the normalized intensity and portality were written to a data frame for subsequent analysis.

### Lobular geometries

For every detected lobulus the following geometric parameters were calculated based on the polygon for the lobulus boundary: perimeter, area, compactness and minimum spanning distance.

### Calculation of number of required lobules

#### Geometric parameters

The number of required lobules n was calculated based on the method of determining sample size for estimating a population mean. The method depends on the margin of error ME (how accurate the results should be), a given confidence level of 95%, i.e. α = 0.05 (how confident the results need to be), and the estimate for the mean and standard deviation.

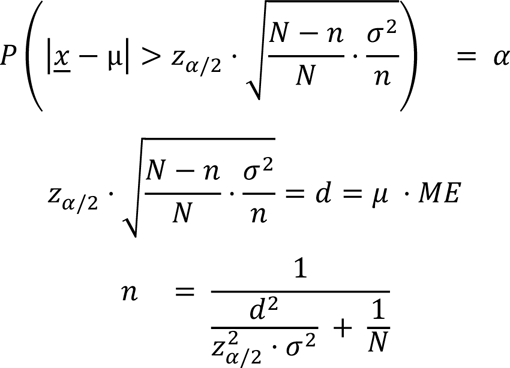

For an analyzed subject and a geometric parameter, N is the total number of lobules, μ is the mean of the parameter over all lobuli of a subject, σ the corresponding standard deviation (provided in Supplementary Table 1), d the distance to the mean based on the margin of error ME, and z the z statistic for a given α value.

#### Zonation patterns

The number of lobuli n required for the zonation patterns was calculated analogously to the geometric parameters. For each subject and each protein, one n was determined for each of the 12 zones. The mean µ is the mean of the intensity for the given position over the lobuli, with σ the corresponding standard deviation and N the number of lobuli for a subject. A margin of error ME of 10% and a confidence level of 95% (α=0.05) were used for the calculation.

#### Relative expression

Relative expression was calculated for each lobule as the sum of all normalized intensities in the lobule for the respective protein. The mean μ and standard deviation σ of relative expression for all lobules of a subject were used to analyze the required number of lobules n, where N is the number of all lobules for a subject. A margin of error ME of 10% and a confidence level of 95% (α=0.05) were used for the calculation.

## Author contribution

MA, MK and UD designed the study and drafted the manuscript. MA selected the samples from the archive, established and performed the staining, performed qualitative and quantitative analysis of all samples. JK and MK developed, implemented and applied the pipeline for image analysis and performed the quantification of lobulus geometry, zonated quantification, and number of required lobulus and created the respective figures and analyses. JG supported the image analysis. OD supervised the histological analysis, discussed the results and reviewed the manuscript, HMT designed and implemented the biobanking of clinical samples and contributed to the study design, US, SS and RK critically discussed the manuscript, EKI supported the human sample collection, VM contributed the pig samples, NN supported the collection of pig samples. All authors read and approved the manuscript.

## Supplementary figures & tables

**Supplementary Figure 1.**
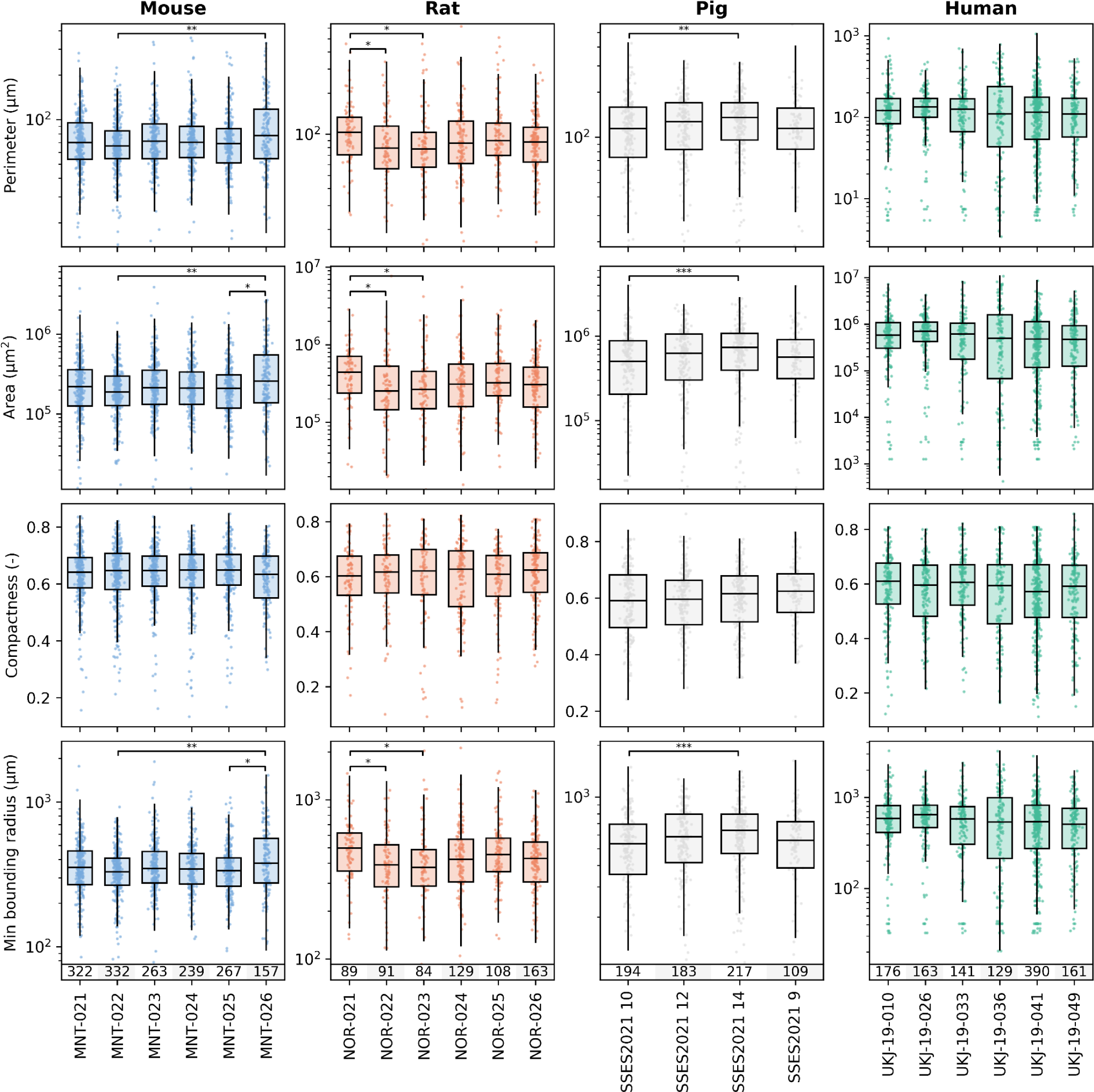
Intra- and inter-individual variability of lobular geometric parameters in human, pig, rat and mouse. Boxes represent quantiles Q1 and Q3. Upper and lower whiskers extend to the last date less than Q3 + 1.5 * IQR and the first date greater than Q1 - 1.5 * IQR, respectively. IQR denotes interquartile range (Q3-Q1). Significance levels: * p<0.05, ** p<0.01, *** p<0.001, **** p<0.0001.

**Supplementary Figure 2.**
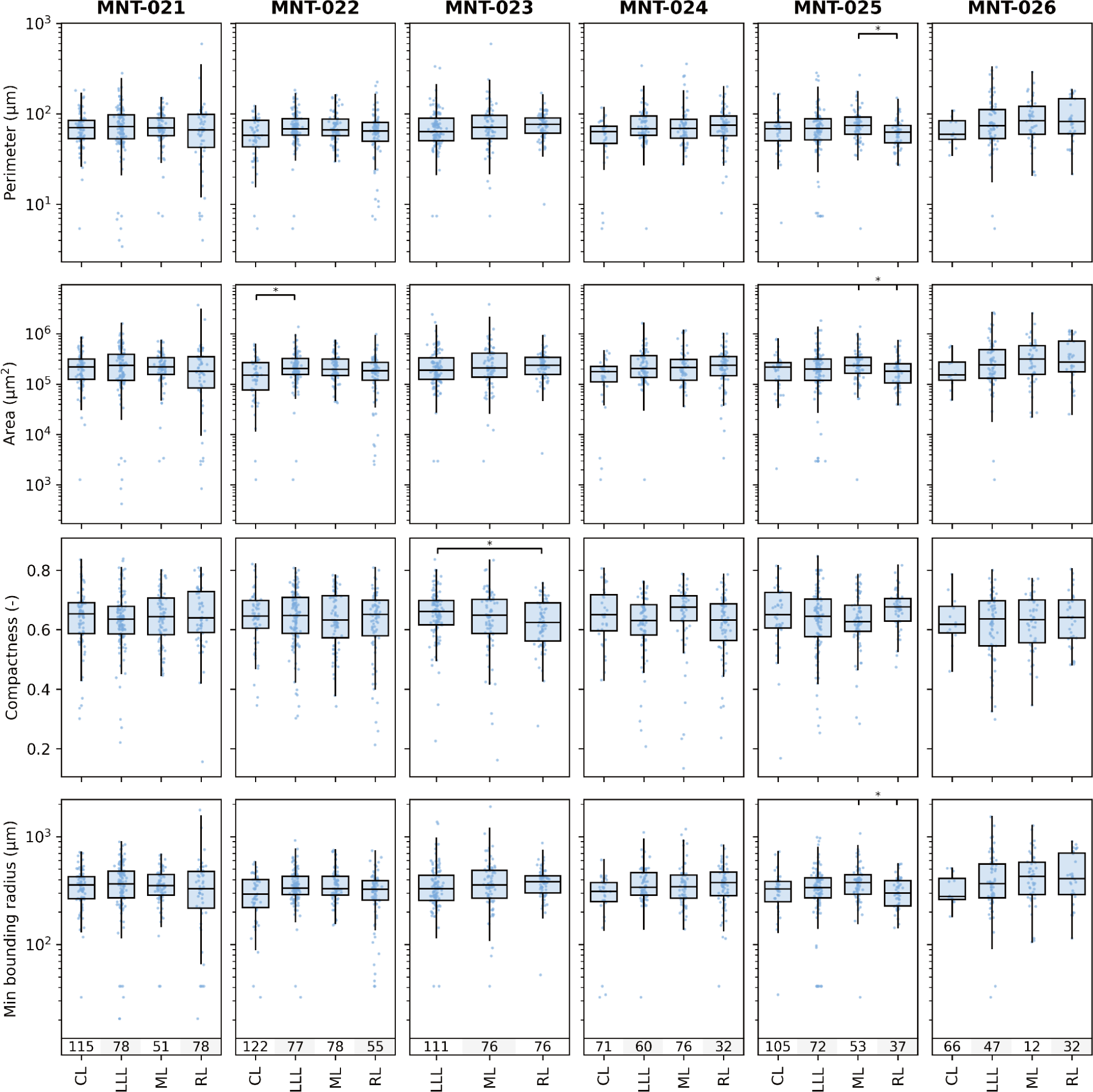
Intra-lobe variability in lobular geometric parameters in mice. Boxes represent quantiles Q1 and Q3. Upper and lower whiskers extend to the last date less than Q3 + 1.5 * IQR and the first date greater than Q1 - 1.5 * IQR, respectively. IQR denotes interquartile range (Q3-Q1). Significance levels: * p<0.05, ** p<0.01, *** p<0.001, **** p<0.0001. CL: caudate lobe, LLL: left lateral lobe, ML: median lobe, right lobe; lCL in MNT-023 could not be evaluated due to lack of ROI registration.

**Supplementary Table 1.** Overview of geometric parameters between species.

| species | parameter | n | mean | sd | se | median | min | max | q1 | q3 | unit |
| --- | --- | --- | --- | --- | --- | --- | --- | --- | --- | --- | --- |
| mouse | perimeter | 1580 | 76.1 | 39.6 | 1.9 | 69.6 | 3.4 | 450.2 | 53.9 | 90.9 | µm |
| rat | perimeter | 664 | 96.1 | 62.4 | 3.7 | 86.5 | 5.4 | 685.1 | 62.8 | 116.8 | µm |
| pig | perimeter | 703 | 121.1 | 60.6 | 4.6 | 122.6 | 5.4 | 446.8 | 82.9 | 158.0 | µm |
| human | perimeter | 1160 | 130.9 | 98.9 | 3.8 | 117.0 | 3.4 | 963.5 | 64.8 | 170.0 | µm |
| mouse | area | 1580 | 282730 | 292549 | 7113 | 210042 | 423 | 4079840 | 128500 | 339493 | µm<sup>2</sup> |
| rat | area | 664 | 470198 | 646351 | 18247 | 305861 | 1269 | 8682985 | 171333 | 558207 | µm<sup>2</sup> |
| pig | area | 703 | 714159 | 565512 | 26935 | 629067 | 1269 | 4071802 | 306073 | 1011499 | µm<sup>2</sup> |
| human | area | 1160 | 879809 | 1159576 | 25832 | 556303 | 423 | 10847280 | 174189 | 1125827 | µm<sup>2</sup> |
| mouse | compactness | 1580 | 0.64 | 0.09 | 0.02 | 0.65 | 0.13 | 0.85 | 0.59 | 0.70 | - |
| rat | compactness | 664 | 0.62 | 0.10 | 0.02 | 0.63 | 0.23 | 0.83 | 0.57 | 0.69 | - |
| pig | compactness | 703 | 0.62 | 0.09 | 0.02 | 0.63 | 0.28 | 0.90 | 0.56 | 0.68 | - |
| human | compactness | 1160 | 0.59 | 0.12 | 0.02 | 0.61 | 0.14 | 0.86 | 0.52 | 0.67 | - |
| mouse | minimum_bounding_radius | 1580 | 372.6 | 178.6 | 9.4 | 344.6 | 20.6 | 1903.2 | 269.1 | 442.36 | µm |
| rat | minimum_bounding_radius | 664 | 463.8 | 270.2 | 18.0 | 424.2 | 32.5 | 2461.1 | 314.0 | 559.71 | µm |
| pig | minimum_bounding_radius | 703 | 580.4 | 276.0 | 21.9 | 584.1 | 32.5 | 1629.0 | 409.6 | 759.00 | µm |
| human | minimum_bounding_radius | 1160 | 615.1 | 419.9 | 18.1 | 572.6 | 20.6 | 3258.9 | 309.4 | 820.02 | µm |

**Supplementary Table 2.** Overview of required lobuli to determine geometric parameters in different species (with 95% confidence and a 10% margin of error).

| species | Perimeter (µm) | Area (µm<sup>2</sup>) | compactness | Minimum_bounding_radius (µm) |
| --- | --- | --- | --- | --- |
| human | 94.9 ± 24.8 | 136 ± 43.4 | 13.4 ± 1.9 | 85.7 ± 18.0 |
| pig | 61.2 ± 6.6 | 99.3 ± 13.4 | 8.2 ± 1.0 | 57.0 ± 6.7 |
| rat | 61.9 ± 9.7 | 89.7 ± 15.2 | 9.3 ± 1.1 | 57.0 ± 9.2 |
| mouse | 70.3 ± 9.5 | 146.6 ± 25.8 | 7.7 ± 1.0 | 62.5 ± 8.2 |

**Supplementary Table 3.** Overview of required lobuli to determine geometric parameters in different subjects.

| species | subject | parameter | unit | mean | std | n | n0.05 | n0.1 | n0.15 | n0.2 | n0.25 | n0.3 | n0.35 |
| --- | --- | --- | --- | --- | --- | --- | --- | --- | --- | --- | --- | --- | --- |
| human | UKJ-19-010_Human | perimeter | µm | 136.8 | 100.2 | 176 | 145.0 | 94.9 | 60.3 | 39.9 | 27.8 | 20.3 | 15.4 |
| human | UKJ-19-026_Human | perimeter | µm | 134.1 | 67.8 | 163 | 115.3 | 61.3 | 34.5 | 21.4 | 14.4 | 10.2 | 7.7 |
| human | UKJ-19-033_Human | perimeter | µm | 130.7 | 98.7 | 141 | 121.4 | 85.7 | 57.6 | 39.4 | 28.0 | 20.7 | 15.9 |
| human | UKJ-19-036_Human | perimeter | µm | 149.3 | 139.5 | 129 | 117.7 | 93.1 | 69.1 | 50.8 | 37.9 | 28.9 | 22.6 |
| human | UKJ-19-041_Human | perimeter | µm | 123.8 | 95.7 | 390 | 273.7 | 144.4 | 80.8 | 50.0 | 33.5 | 23.9 | 17.9 |
| human | UKJ-19-049_Human | perimeter | µm | 123.6 | 90.1 | 161 | 134.5 | 90.0 | 58.0 | 38.8 | 27.2 | 19.9 | 15.1 |
| pig | SSES2021 10 | perimeter | µm | 111.1 | 61.0 | 194 | 136.8 | 72.6 | 40.7 | 25.2 | 16.9 | 12.1 | 9.0 |
| pig | SSES2021 12 | perimeter | µm | 123.8 | 57.7 | 183 | 118.2 | 57.3 | 30.8 | 18.7 | 12.4 | 8.8 | 6.6 |
| pig | SSES2021 14 | perimeter | µm | 130.4 | 58.9 | 217 | 128.3 | 57.6 | 30.1 | 18.0 | 11.9 | 8.4 | 6.2 |
| pig | SSES2021 9 | perimeter | µm | 115.9 | 64.8 | 109 | 88.8 | 57.1 | 35.8 | 23.5 | 16.3 | 11.9 | 9.0 |
| rat | NOR-021 | perimeter | µm | 106.9 | 56.6 | 89 | 73.8 | 48.7 | 31.1 | 20.7 | 14.4 | 10.5 | 8.0 |
| rat | NOR-022 | perimeter | µm | 90.8 | 77.9 | 91 | 84.2 | 68.8 | 52.8 | 39.8 | 30.2 | 23.3 | 18.4 |
| rat | NOR-023 | perimeter | µm | 93.2 | 70.9 | 84 | 76.7 | 60.9 | 45.4 | 33.4 | 25.0 | 19.1 | 14.9 |
| rat | NOR-024 | perimeter | µm | 97.1 | 70.7 | 129 | 111.4 | 79.0 | 53.2 | 36.5 | 26.0 | 19.3 | 14.7 |
| rat | NOR-025 | perimeter | µm | 103.8 | 58.2 | 108 | 88.3 | 57.0 | 35.8 | 23.6 | 16.4 | 11.9 | 9.0 |
| rat | NOR-026 | perimeter | µm | 88.9 | 42.3 | 163 | 110.9 | 56.6 | 31.2 | 19.1 | 12.8 | 9.1 | 6.8 |
| mouse | MNT-021 | perimeter | µm | 75.6 | 40.6 | 322 | 186.3 | 82.3 | 42.6 | 25.4 | 16.8 | 11.8 | 8.8 |
| mouse | MNT-022 | perimeter | µm | 69.9 | 28.5 | 332 | 144.4 | 53.6 | 26.2 | 15.2 | 9.9 | 7.0 | 5.1 |
| mouse | MNT-023 | perimeter | µm | 77.9 | 42.0 | 263 | 165.4 | 78.3 | 41.7 | 25.2 | 16.7 | 11.8 | 8.8 |
| mouse | MNT-024 | perimeter | µm | 76.7 | 38.8 | 239 | 148.7 | 69.7 | 37.0 | 22.3 | 14.8 | 10.5 | 7.8 |
| mouse | MNT-025 | perimeter | µm | 72.6 | 33.9 | 267 | 148.6 | 63.8 | 32.7 | 19.4 | 12.8 | 9.0 | 6.7 |
| mouse | MNT-026 | perimeter | µm | 92.5 | 56.1 | 157 | 122.9 | 74.4 | 44.9 | 28.9 | 19.8 | 14.3 | 10.8 |
| human | UKJ-19-010_Human | area | µm<sup>2</sup> | 905036 | 1106446 | 176 | 163.5 | 134.7 | 104.2 | 79.1 | 60.4 | 46.8 | 37.0 |
| human | UKJ-19-026_Human | area | μm<sup>2</sup> | 833735 | 681768 | 163 | 140.7 | 99.7 | 67.1 | 46.1 | 32.8 | 24.3 | 18.6 |
| human | UKJ-19-033_Human | area | μm<sup>2</sup> | 907595 | 1249543 | 141 | 134.5 | 118.1 | 98.2 | 79.5 | 63.8 | 51.4 | 41.8 |
| human | UKJ-19-036_Human | area | μm<sup>2</sup> | 1373609 | 2058582 | 129 | 124.4 | 112.2 | 96.5 | 80.7 | 66.7 | 55.0 | 45.6 |
| human | UKJ-19-041_Human | area | μm<sup>2</sup> | 761874 | 923075 | 390 | 332.5 | 230.6 | 152.6 | 103.5 | 73.3 | 54.0 | 41.2 |
| human | UKJ-19-049_Human | area | μm<sup>2</sup> | 764575 | 897943 | 161 | 149.6 | 123.5 | 95.6 | 72.7 | 55.5 | 43.1 | 34.1 |
| pig | SSES2021 10 | area | μm<sup>2</sup> | 613730 | 536179 | 194 | 166.5 | 116.8 | 78.0 | 53.2 | 37.8 | 27.9 | 21.3 |
| pig | SSES2021 12 | area | μm<sup>2</sup> | 719448 | 529996 | 183 | 150.1 | 97.5 | 61.5 | 40.6 | 28.2 | 20.6 | 15.6 |
| pig | SSES2021 14 | area | μm<sup>2</sup> | 819219 | 589013 | 217 | 170.4 | 103.7 | 62.7 | 40.4 | 27.7 | 20.0 | 15.1 |
| pig | SSES2021 9 | area | μm<sup>2</sup> | 674868 | 589161 | 109 | 99.7 | 79.4 | 59.3 | 43.8 | 32.8 | 25.1 | 19.6 |
| rat | NOR-021 | area | μm<sup>2</sup> | 544853 | 489037 | 89 | 83.0 | 69.1 | 54.0 | 41.4 | 31.8 | 24.8 | 19.7 |
| rat | NOR-022 | area | μm<sup>2</sup> | 467757 | 954073 | 91 | 89.7 | 86.1 | 80.7 | 74.1 | 67.1 | 60.2 | 53.6 |
| rat | NOR-023 | area | μm<sup>2</sup> | 468567 | 749160 | 84 | 82.2 | 77.4 | 70.4 | 62.6 | 54.7 | 47.5 | 41.0 |
| rat | NOR-024 | area | μm<sup>2</sup> | 485789 | 749686 | 129 | 124.6 | 113.1 | 97.9 | 82.5 | 68.6 | 56.9 | 47.3 |
| rat | NOR-025 | area | μm<sup>2</sup> | 510552 | 547947 | 108 | 101.8 | 86.8 | 69.7 | 54.6 | 42.8 | 33.8 | 27.1 |
| rat | NOR-026 | area | μm<sup>2</sup> | 392562 | 346075 | 163 | 143.4 | 105.4 | 73.1 | 51.2 | 36.9 | 27.6 | 21.2 |
| mouse | MNT-021 | area | μm<sup>2</sup> | 283236 | 304336 | 322 | 272.5 | 186.6 | 122.3 | 82.5 | 58.1 | 42.7 | 32.5 |
| mouse | MNT-022 | area | μm<sup>2</sup> | 232120 | 171162 | 332 | 237.6 | 128.2 | 72.5 | 45.1 | 30.4 | 21.7 | 16.2 |
| mouse | MNT-023 | area | μm<sup>2</sup> | 296800 | 347930 | 263 | 233.9 | 175.5 | 124.0 | 87.9 | 63.9 | 48.0 | 37.0 |
| mouse | MNT-024 | area | μm<sup>2</sup> | 274317 | 234388 | 239 | 197.0 | 129.0 | 81.9 | 54.2 | 37.8 | 27.6 | 20.9 |
| mouse | MNT-025 | area | μm<sup>2</sup> | 260209 | 233906 | 267 | 219.7 | 143.5 | 91.0 | 60.1 | 41.9 | 30.5 | 23.1 |
| mouse | MNT-026 | area | μm<sup>2</sup> | 416249 | 453187 | 157 | 144.5 | 116.7 | 88.4 | 66.0 | 49.8 | 38.3 | 30.1 |
| human | UKJ-19-010_Human | compactness | - | 0.60 | 0.12 | 176 | 43.3 | 13.3 | 6.2 | 3.5 | 2.3 | 1.6 | 1.2 |
| human | UKJ-19-026_Human | compactness | - | 0.60 | 0.10 | 163 | 35.7 | 10.7 | 4.9 | 2.8 | 1.8 | 1.3 | 0.9 |
| human | UKJ-19-033_Human | compactness | - | 0.61 | 0.11 | 141 | 36.9 | 11.5 | 5.3 | 3.1 | 2.0 | 1.4 | 1.0 |
| human | UKJ-19-036_Human | compactness | - | 0.60 | 0.12 | 129 | 41.5 | 13.7 | 6.5 | 3.7 | 2.4 | 1.7 | 1.2 |
| human | UKJ-19-041_Human | compactness | - | 0.58 | 0.12 | 390 | 57.7 | 16.2 | 7.4 | 4.2 | 2.7 | 1.9 | 1.4 |
| human | UKJ-19-049_Human | compactness | - | 0.59 | 0.12 | 161 | 46.7 | 14.9 | 7.0 | 4.0 | 2.6 | 1.8 | 1.3 |
| pig | SSES2021 10 | compactness | - | 0.61 | 0.10 | 194 | 34.3 | 9.9 | 4.5 | 2.6 | 1.7 | 1.2 | 0.8 |
| pig | SSES2021 12 | compactness | - | 0.61 | 0.09 | 183 | 28.8 | 8.2 | 3.7 | 2.1 | 1.4 | 0.9 | 0.7 |
| pig | SSES2021 14 | compactness | - | 0.63 | 0.09 | 217 | 25.5 | 7.0 | 3.2 | 1.8 | 1.1 | 0.8 | 0.6 |
| pig | SSES2021 9 | compactness | - | 0.63 | 0.09 | 109 | 26.0 | 7.9 | 3.7 | 2.1 | 1.4 | 0.9 | 0.7 |
| rat | NOR-021 | compactness | - | 0.62 | 0.10 | 89 | 28.6 | 9.4 | 4.5 | 2.6 | 1.7 | 1.2 | 0.9 |
| rat | NOR-022 | compactness | - | 0.63 | 0.09 | 91 | 24.8 | 7.8 | 3.6 | 2.1 | 1.3 | 0.9 | 0.7 |
| rat | NOR-023 | compactness | - | 0.62 | 0.11 | 84 | 31.1 | 10.8 | 5.2 | 3.0 | 1.9 | 1.4 | 1.0 |
| rat | NOR-024 | compactness | - | 0.62 | 0.11 | 129 | 34.4 | 10.8 | 5.0 | 2.9 | 1.9 | 1.3 | 1.0 |
| rat | NOR-025 | compactness | - | 0.61 | 0.09 | 108 | 27.1 | 8.3 | 3.9 | 2.2 | 1.4 | 1.0 | 0.7 |
| rat | NOR-026 | compactness | - | 0.63 | 0.10 | 163 | 30.6 | 8.9 | 4.1 | 2.3 | 1.5 | 1.0 | 0.8 |
| mouse | MNT-021 | compactness | - | 0.64 | 0.09 | 322 | 27.8 | 7.4 | 3.4 | 1.9 | 1.2 | 0.8 | 0.6 |
| mouse | MNT-022 | compactness | - | 0.64 | 0.09 | 332 | 28.3 | 7.6 | 3.4 | 1.9 | 1.2 | 0.9 | 0.6 |
| mouse | MNT-023 | compactness | - | 0.64 | 0.08 | 263 | 24.4 | 6.6 | 3.0 | 1.7 | 1.1 | 0.7 | 0.5 |
| mouse | MNT-024 | compactness | - | 0.63 | 0.10 | 239 | 31.6 | 8.8 | 4.0 | 2.3 | 1.4 | 1.0 | 0.7 |
| mouse | MNT-025 | compactness | - | 0.65 | 0.09 | 267 | 24.3 | 6.5 | 2.9 | 1.7 | 1.1 | 0.7 | 0.5 |
| mouse | MNT-026 | compactness | - | 0.62 | 0.10 | 157 | 31.5 | 9.3 | 4.3 | 2.4 | 1.6 | 1.1 | 0.8 |
| human | UKJ-19-010_Human | minimum_bounding_radius | μm | 653.2 | 415.8 | 176 | 137.2 | 82.6 | 49.7 | 31.9 | 21.8 | 15.8 | 11.9 |
| human | UKJ-19-026_Human | minimum_bounding_radius | μm | 648.8 | 315.9 | 163 | 112.6 | 58.4 | 32.4 | 20.0 | 13.4 | 9.5 | 7.1 |
| human | UKJ-19-033_Human | minimum_bounding_radius | μm | 619.1 | 424.5 | 141 | 118.0 | 79.2 | 51.2 | 34.2 | 24.0 | 17.6 | 13.4 |
| human | UKJ-19-036_Human | minimum_bounding_radius | μm | 688.4 | 611.7 | 129 | 116.6 | 90.5 | 65.9 | 47.8 | 35.3 | 26.7 | 20.8 |
| human | UKJ-19-041_Human | minimum_bounding_radius | μm | 572.1 | 382.5 | 390 | 248.8 | 119.2 | 63.8 | 38.7 | 25.7 | 18.2 | 13.5 |
| human | UKJ-19-049_Human | minimum_bounding_radius | μm | 581.3 | 394.4 | 161 | 131.2 | 84.3 | 52.8 | 34.7 | 24.1 | 17.5 | 13.3 |
| pig | SSES2021 10 | minimum_bounding_radius | μm | 527.9 | 277.4 | 194 | 133.1 | 68.6 | 37.9 | 23.3 | 15.6 | 11.1 | 8.3 |
| pig | SSES2021 12 | minimum_bounding_radius | μm | 590.7 | 265.0 | 183 | 115.0 | 54.3 | 28.9 | 17.5 | 11.6 | 8.2 | 6.1 |
| pig | SSES2021 14 | minimum_bounding_radius | µm | 630.1 | 271.3 | 217 | 123.2 | 53.6 | 27.6 | 16.4 | 10.8 | 7.6 | 5.7 |
| pig | SSES2021 9 | minimum_bounding_radius | µm | 557.6 | 282.0 | 109 | 85.3 | 51.7 | 31.2 | 20.0 | 13.7 | 9.9 | 7.5 |
| rat | NOR-021 | minimum_bounding_radius | µm | 515.0 | 247.5 | 89 | 71.2 | 44.4 | 27.3 | 17.8 | 12.2 | 8.9 | 6.7 |
| rat | NOR-022 | minimum_bounding_radius | µm | 436.6 | 307.8 | 91 | 81.3 | 61.6 | 43.9 | 31.3 | 22.9 | 17.2 | 13.3 |
| rat | NOR-023 | minimum_bounding_radius | µm | 443.4 | 302.3 | 84 | 75.2 | 57.1 | 40.8 | 29.2 | 21.3 | 16.1 | 12.4 |
| rat | NOR-024 | minimum_bounding_radius | µm | 468.3 | 314.2 | 129 | 108.7 | 73.9 | 48.2 | 32.4 | 22.8 | 16.7 | 12.7 |
| rat | NOR-025 | minimum_bounding_radius | µm | 503.3 | 254.1 | 108 | 84.7 | 51.3 | 31.0 | 20.0 | 13.7 | 9.9 | 7.4 |
| rat | NOR-026 | minimum_bounding_radius | µm | 431.7 | 197.4 | 163 | 108.1 | 53.8 | 29.3 | 17.9 | 11.9 | 8.5 | 6.3 |
| mouse | MNT-021 | minimum_bounding_radius | µm | 372.0 | 181.5 | 322 | 171.2 | 71.2 | 36.1 | 21.3 | 14.0 | 9.8 | 7.3 |
| mouse | MNT-022 | minimum_bounding_radius | µm | 343.7 | 134.1 | 332 | 137.3 | 49.7 | 24.1 | 14.0 | 9.1 | 6.4 | 4.7 |
| mouse | MNT-023 | minimum_bounding_radius | µm | 385.6 | 195.0 | 263 | 157.6 | 71.5 | 37.5 | 22.5 | 14.8 | 10.5 | 7.8 |
| mouse | MNT-024 | minimum_bounding_radius | µm | 372.5 | 165.0 | 239 | 133.3 | 57.3 | 29.4 | 17.5 | 11.5 | 8.1 | 6.0 |
| mouse | MNT-025 | minimum_bounding_radius | µm | 354.0 | 154.5 | 267 | 139.6 | 57.4 | 29.0 | 17.1 | 11.2 | 7.9 | 5.8 |
| mouse | MNT-026 | minimum_bounding_radius | µm | 445.2 | 248.8 | 157 | 118.3 | 68.0 | 39.8 | 25.2 | 17.1 | 12.3 | 9.2 |

**Supplementary Table 4.** Overview of required lobuli to determine Relative GS and CYPs expression in different species (with 95% confidence and a 10% margin of error).

| species | HE | CYP1A2 | CYP2D6 | CYP2E1 | CYP3A4 |
| --- | --- | --- | --- | --- | --- |
| human | 9.5 ± 2.1 | 23.1 ± 6.4 | 15.2 ± 7.6 | 18.2 ± 4.0 | 27.7 ± 11.3 |
| pig | 10.2 ± 6.9 | 16.9 ± 5.8 | 15.4 ± 3.2 | 14.9 ± 2.7 | 10.5 ± 3.5 |
| rat | 8.0 ± 6.6 | 5.8 ± 3.3 | 6.4 ± 0.8 | 5.8 ± 2.0 | 4.5 ± 1.9 |
| mouse | 4.3 ± 2.8 | 6.3 ± 2.7 | 3.6 ± 0.9 | 6.0 ± 1.8 | 6.9 ± 2.0 |

**Supplementary Table 5.** Antibodies used for IHC-staining of GS and CYP enzymes for qualitative evaluation of expression pattern and signal intensity and quantitation of spatial distribution and zonation (m for mouse, R for rat, p for pig, and h for human)

| Antibody | Company | Order-Nr | Dilution/species | Detection systems |
| --- | --- | --- | --- | --- |
| Anti-CYP2D6 (Rabbit polyclonal antibody) | Abcam, Germany | ab230690 | 1/3000 (m), 13000 (R), 1/2000 (p), 1/200 (H) | Rabbit-specific HRP/DAB Detection IHC Detection Kit - Micro-polymer (ab236469, Abcam, Germany) |
| Anti-CYP2E1 (Rabbit polyclonal antibody) | Sigma-Aldrich, Germany | HPA009128 | 1/400 (m), 1/300 R, 1/800 p, 1/800 h |  |
| Anti-CYP3A4 (Rabbit polyclonal antibody) | Abcam, Germany | ab3572 | 1/2000 (m), 1/2000 R, 1/1000 p, 1/1500 h |  |
| Anti-CYP1A2 (Mouse monoclonal antibody) | Abcam, Germany | ab22717 | 1/500 (m), 1/200 R, 1/2000 p, 1/2000 h | Dako Animal Research Kit Peroxidase for Mouse primary antibody, (K3954, Dako, Denmark) |
| Anti-GS (Mouse monoclonal antibody) | Merck | MAB302 | 1/1000 (m), 1/1000R, 1000 p, 1/1000 h |  |

